# The transcriptional co-repressor CtBP is a negative regulator of growth that antagonizes the Yorkie and JNK/AP-1 pathways

**DOI:** 10.1101/772533

**Authors:** Taryn M. Sumabat, Melanie I. Worley, Brett J. Pellock, Justin A. Bosch, Iswar K. Hariharan

**Affiliations:** Department of Molecular and Cell Biology, University of California, Berkeley, Berkeley, CA 94720-3200; Department of Biology, Providence College, Providence, RI 02918

## Abstract

Multicellular organisms require strict growth control mechanisms to ensure that an organ reaches, but does not grossly exceed, its appropriate size and shape. In an unbiased mosaic screen for genes involved in growth regulation, we identified a loss-of-function allele of the gene *CtBP* that conferred a growth advantage to homozygous mutant tissue. *CtBP* encodes a transcriptional co-repressor found in diverse organisms, yet its role in regulating tissue growth is not known. We found that *CtBP* functions as a negative regulator of growth by restricting the expression of the growth-promoting microRNA *bantam* (*ban*). *ban* is a known target of the Hippo pathway effector Yorkie (Yki). We show that loss of *CtBP* function leads to the activation of a minimal enhancer of *ban* via both Yki-dependent and AP-1 transcription factor-dependent mechanisms. AP-1 is downstream of the Jun N-terminal Kinase (JNK) pathway and thus JNK could regulate growth during development via *ban*. Furthermore, we show that distinct isoforms of the AP-1 component Fos differ in their ability to activate this enhancer. Since the orthologous pathways in mammalian cells (YAP/TEAD and AP-1) converge on enhancers implicated in tumor progression, a role for mammalian CtBP proteins at those enhancers merits attention.

## Introduction

Tissue growth is a fundamental aspect of animal development. Strict control mechanisms are needed to ensure that an organ does not exceed its correct size at the end of development, and to allow for regenerative growth upon tissue damage. Although individual cells are the generators of biomass, organs composed of thousands of cells are able to stop growing when they reach a predictable size and shape. The mechanisms that link the size and shape of the organ to the proliferation of thousands of individual cells are still poorly understood.

The Hippo pathway, initially discovered in *Drosophila*, is essential for normal development and plays a key role in growth control, through the coordinated regulation of cell proliferation and apoptosis (recently reviewed by (Misra and Irvine, 2018; Zheng and Pan, 2019)). Many of the pathway components found in *Drosophila* are shared among diverse metazoans (Sebe-Pedros *et al.*, 2012), with great functional conservation in mammals; (Zhao *et al.*, 2011). The *Drosophila* Hippo pathway restricts tissue growth by inhibiting the nuclear localization of the transcriptional coactivator Yorkie (Yki) (Huang *et al.*, 2005) Yki does not bind DNA itself, but is targeted to specific loci in the genome via its association with sequence-specific DNA binding proteins, most notably Scalloped (Sd) (Wu *et al.*, 2008; Zhang *et al.*, 2008). Yki has also been proposed to function in complex with other DNA binding proteins as well including Teashirt and Homothorax in the eye disc (Peng *et al.*, 2009) and Mad (Oh and Irvine, 2011) although these claims have been contested (Yu and Pan, 2018). Under pro-growth conditions, Yki can accumulate in the nucleus, where it activates numerous target genes. Among Yki targets is the microRNA *bantam* (*ban*), which is capable of both promoting growth and inhibiting cell death (Nolo *et al.*, 2006; Thompson and Cohen, 2006).

While we are far from a complete understanding of the transcriptional program downstream of Yki that defines its role in developmental growth, mounting evidence suggests that the components of this program are likely subject to multiple regulatory inputs. Indeed, several studies have demonstrated that Yki-dependent regulatory enhancers often contain consensus binding sites for the transcriptional effectors of other conserved signaling pathways (Oh *et al.*, 2013; Pascual *et al.*, 2017). A transcription factor that has been shown to have binding sites in the genetic loci for many Yki targets is AP-1 (Pascual *et al.*, 2017), the downstream transcriptional effector of the c-Jun amino-terminal kinase (JNK) pathway (Eferl and Wagner, 2003). The JNK pathway is a kinase cascade involved in several morphogenetic processes during development and is activated in response to various stress stimuli. JNK signaling has been shown to trigger apoptosis or tumorigenesis, depending on the genetic context, via activation of AP-1. AP-1 are homodimeric and heterodimeric protein complexes consisting of the bZIP-domain containing proteins Fos and Jun. As in *Drosophila*, studies using human cancer cell lines have shown that the human orthologs of Yki, YAP and TAZ, localize to enhancers in complex with the Sd ortholog TEAD4 in proximity to AP-1 binding sites and that TEAD/YAP and AP-1 synergistically regulate the expression of genes that promote cancer progression (Verfaillie *et al.*, 2015; Zanconato *et al.*, 2015; Liu *et al.*, 2016). The mechanisms that regulate synergy between these two pathways are still not well characterized.

We had previously identified *CtBP*, which encodes a transcriptional corepressor, in a screen for genes that regulate regenerative plasticity (Worley *et al.*, 2018). In a separate screen, we identified an allele of *CtBP* as a mutation that enables mutant cells to outgrow their wild-type neighbors. Here we characterize the role of *CtBP* as a negative regulator of growth and show that this is due, in significant part, to its effect on reducing expression of the growth effector *ban*. We show that *CtBP* regulates a minimal *ban* enhancer via both *yki*-dependent and *yki*-independent mechanisms. In addition to *yki*, we show that the JNK/AP-1 pathway can activate this enhancer and that *CtBP* appears capable of antagonizing both of these pathways. Thus, *CtBP* maintains the appropriate level of growth during development by modulating the activity of both the Yki and JNK/AP-1 pathways.

## Materials and Methods

### *Drosophila* stocks and husbandry

Crosses were maintained on standard fly food at 25°C unless otherwise noted. For experiments sensitive to the effects of developmental staging and vial density, egg deposition was limited to a 4 h window.

The *CtBP*^*A147T*^ allele is an EMS mutation induced on an *FRT82B* chromosome. The *CtBP*^*Q229\**^ allele (BDSC 1663) has been described as a null allele and was recombined onto the parental *FRT82B* chromosome for use in this study. The *CtBP*^*N148fs*^ mutation was generated on the *FRT82B* chromosome using CRISPR/Cas9 and a guide RNA targeting codon 148.

Additional stocks used were: *FRT82B* (Xu and Rubin, 1993), *GFP-ban* (*bantam* sensor, (Brennecke *et al.*, 2003)); *ban3-GFP* (Matakatsu and Blair, 2012); *ex*^*697*^ (Boedigheimer and Laughon, 1993); *Diap1 3.5-GFP* (Zhang *et al.*, 2008); *FRT42D yki*^*B5*^ (Huang *et al.*, 2005); *FRT42D MARCM*; *UAS-ban-sponge* (Becam *et al.*, 2011); *UAS-Fos.D* (Ling *et al.*, 2012).

Remaining stocks used in this study were from, or derived from, the Bloomington Stock Center (Bloomington, IN): *FRT82B ubi-RFPnls* (BDSC 30555); *FRT82B MARCM* (BDSC 30036); *UbxFlp; FRT42D GFP*; *FRT82B RFP* (BDSC 43340); *FRT82B kay*^*ED6315*^ (BDSC 41772); *brC12-lacZ* (BDSC 44256); *puc*^*A251.1F3*^ (BDSC 11173); *nub-Gal4* (BDSC 25754); *ptc-Gal4* (BDSC 2017); UAS-GFP (BDSC 6874); UAS-yellow-shRNA (BDSC 64527); *UAS-CtBP-shRNA* (BDSC 32889); *UAS-puc-shRNA* (BDSC 53019); *UAS-Hep-WT* (BDSC 9308); *UAS-Bsk-DN* (BDSC 9311); *UAS-bsk-shRNA* (BDSC 57035); *UAS-Fos* (BDSC 7213); *UAS-fos-shRNA* (BDSC 33379); *UAS-Jun* (BDSC 7216).

### Cloning and molecular biology

Transgenic *UAS-CtBP-HA* stocks for overexpression of each unique CtBP protein isoform were generated for this study and integrated at the *attP40* landing site. We generated a stable transgenic with a CRISPR guide RNA targeting a conserved region in the *CtBP* gene, following the method described in the FlyCas9 system (Kondo and Ueda, 2013).

### Clonal Analysis

For the clone and twin-spot quantification experiments shown in Figure 1, clones were induced 48 h after egg deposition with a 45±15 min heat-shock at 37°C and wing imaginal discs were dissected 72 h later. For an individual mosaic disc, the total clone area was determined by tracing all the RFP-negative clones and adding up their areas, and the total twin-spot area was determined by tracing all the 2 RFP-positive clones and adding up their areas.

**Figure 1.**
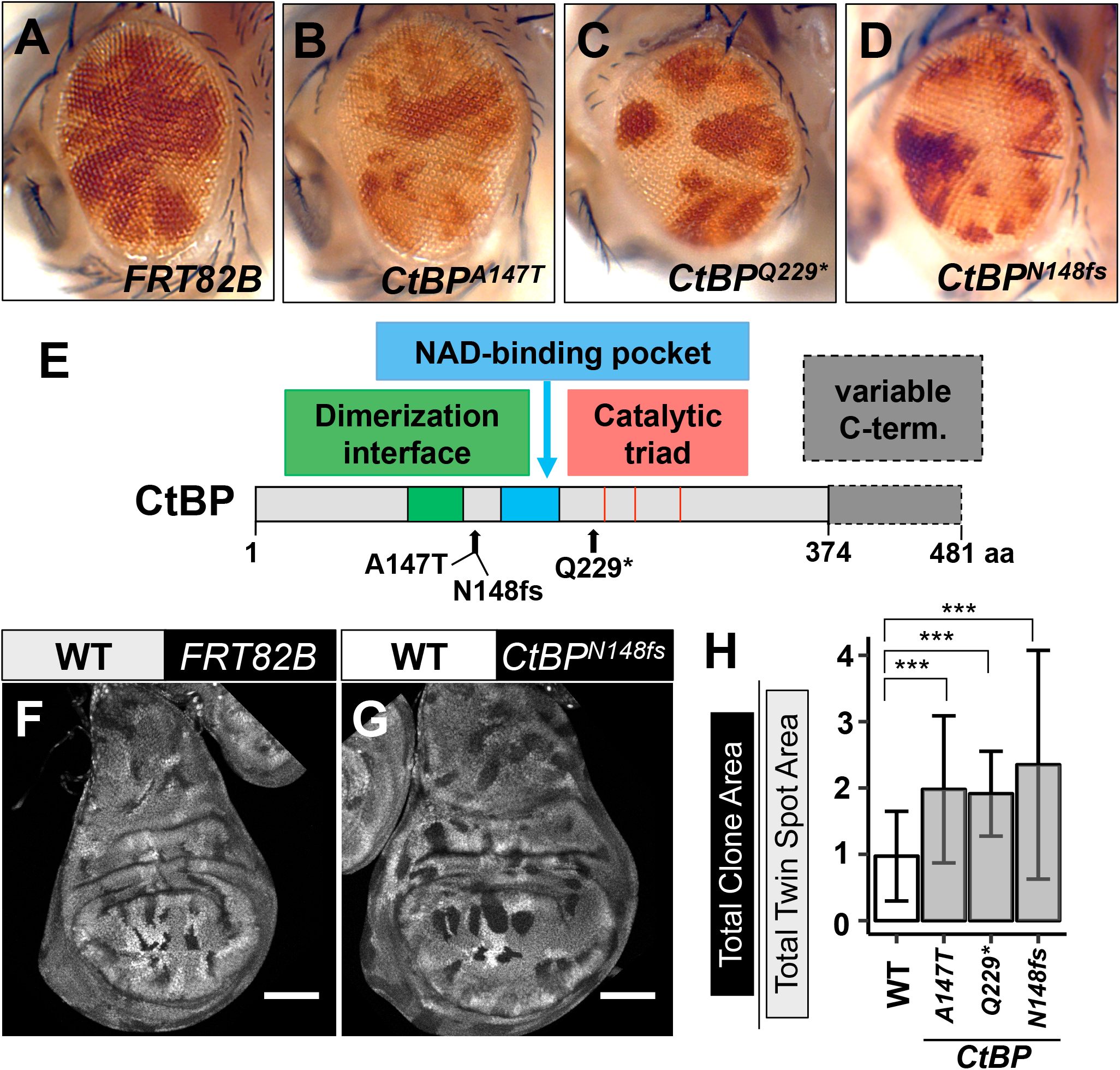
Mutations in *CtBP* result in increased tissue growth. (A-D) Mosaic eyes containing homozygous mutant clones (white) and wild-type twins spots (red). (A) Control mosaic eyes. (B) *CtBP*^*A147T*^, (C) *CtBP*^*Q229\**^, and (D) *CtBP*^*N148fs*^ mosaic eyes have more mutant thn wild-type tissue. (E) Protein model of Drosophila CtBP showing general domain structures and the three coding mutations used in this study. The light gray region is shared by all predicted isoforms. The dark gray region is present only in some isoforms. (F-G) *hsFlp* mosaic Wing imaginal discs containing mutant clones and wild-type twin spots. Mutant tissue is marked by the absence of a fluorescent marker, twin-spot clones contain two copies of the marker. Scale bars, 100 μm. (H) Graph showing the ratio of total mutant tissue to total wild-type twin-spot tissue measured from mosaic wing discs. “Total clone area” was determined by tracing all the unmarked clones from a disc and adding up their areas. “Total twin-spot area” was determined by tracing all the 2xRFP-positive clones from a disc and adding up their areas. n =12, 9, 9, and 5 discs per genotype, respectively. Data are presented as mean ± S.D. Statistical significance was determined using one-way ANOVA with Tukey’s post-hoc test. \*\*\**P* ≤ 0.001.

For the MARCM experiments shown in Figures 2 and 3, clones were induced 48 h after egg deposition with an hour-long 37°C heat-shock, and wing imaginal discs were dissected 72 h later. Individual clones were traced using the Polygon tool and measured in ImageJ.

**Figure 2.**
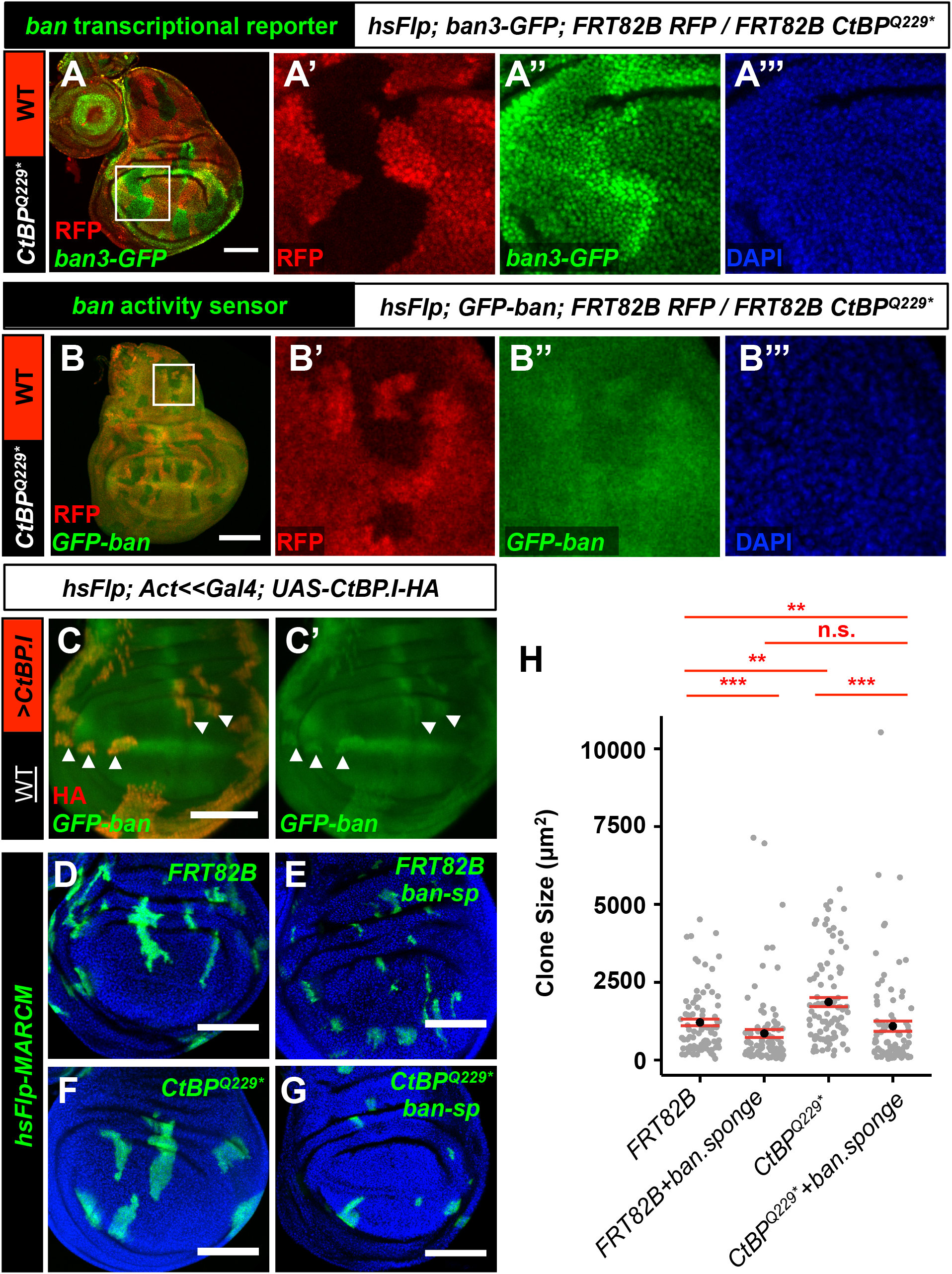
Increased *bantam* activity is required for overgrowth of *CtBP* mutant clones. (A) Mosaic wing imaginal discs with *ban3-GFP* reporter. Wild-type cells are marked with RFP (red) and *CtBP*^*Q229\**^ clones are unmarked. Mutant clones show increased levels of *ban3-GFP* especially near the clone boundary. (B-C) Analysis of the *GFP-ban* sensor (green), which inversely reports *ban* microRNA activity, in clones mutant for or overexpressing *CtBP*. (B-B’’’) *CtBP*^*Q229\**^ clones (unmarked) have lower levels of the *GFP-ban* sensor. (C-C’) Flp-out Gal4 clones overexpressing CtBP show higher levels of the *GFP-ban* sensor. Use of the short isoform CtBP.I-HA is shown and the clones were detected by HA staining (red). Boxes in (A) and (B) indicate areas enlarged in subsequent panels. (D-H) MARCM clone size assay. MARCM Gal4 clones (green) were induced by expression of *hsFlp* following a 1-hour heat-shock at 48 hours after egg-lay. Discs were dissected 72 hours after clone induction (ACI). Shown are representative wing pouches with (D) *FRT82B* control clones, (E) *FRT82B* control clones overexpressing a *ban sponge* (*ban-sp*), (F) *CtBP*^*Q229*\*^ mutant clones, and (G) *CtBP*^*Q229*\*^ mutant clones overexpressing *ban-sp*. (H) Quantification of individual clone sizes (in pixels) from the genotypes shown in D (n=87), E (n=97), F (n=92), and G (n=90), respectively. All data points (gray dots) are shown along with mean (black dots) and SEM (red error bars). Statistical significance was taken as *P* ≤ 0.05 and assessed by performing pairwise comparisons of each MARCM condition using a Wilcoxon rank sum test. *** *P* ≤ 0.001. ** *P* ≤ 0.01. All scale bars in figure, 100 μm. DAPI shows nuclei (blue).

**Figure 3.**
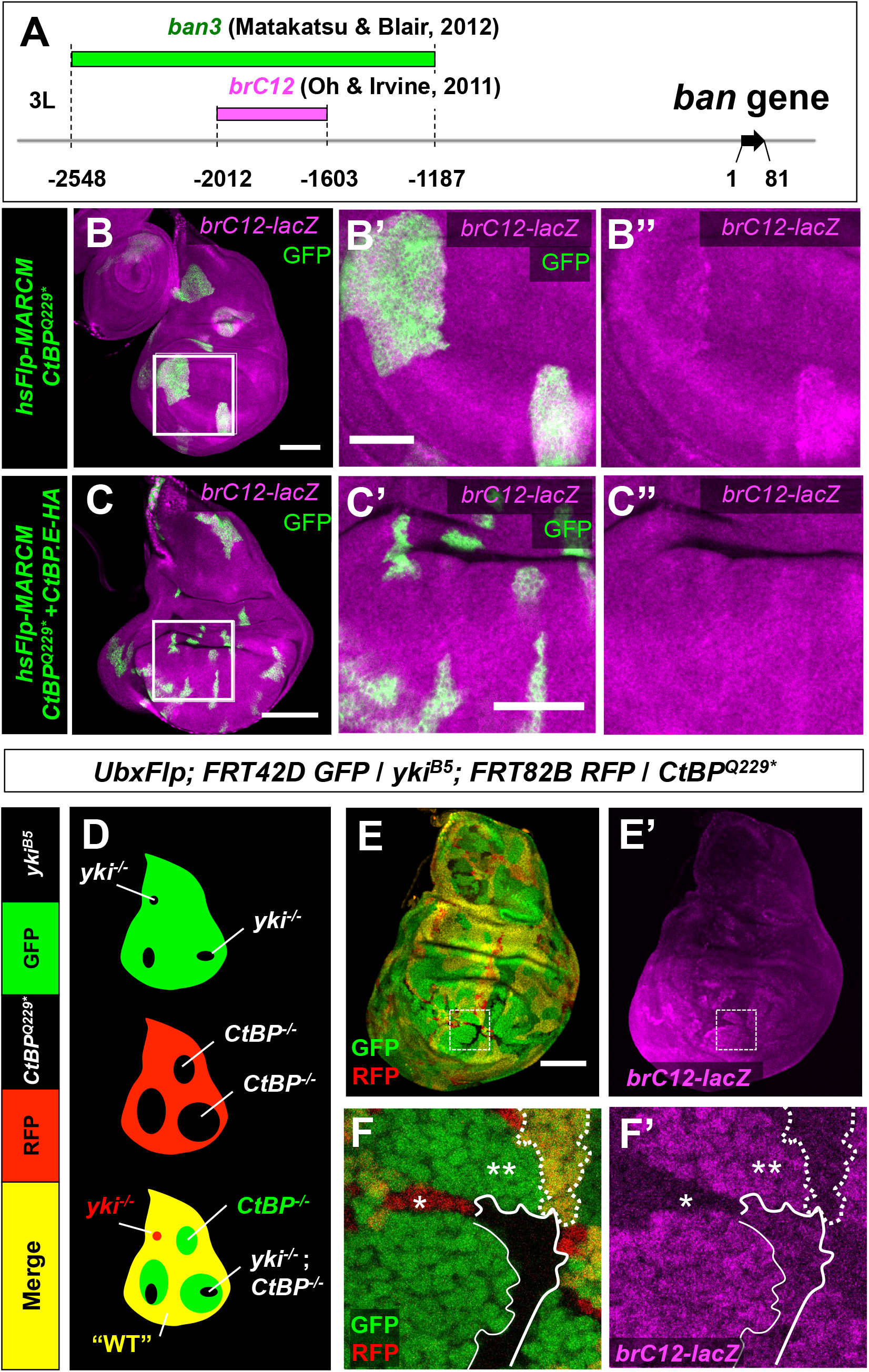
CtBP regulates a minimal *bantam* enhancer via Yki-dependent and Yki-Independent mechanisms. (A) A schematic of the *ban* locus showing the enhancer regions that were cloned into the *ban3-GFP* (green) and *brC12-lacZ* (magenta) reporter transgenes. (B-B”) *CtBP*^*Q229\**^ MARCM clones (green) show increased expression of a minimal *ban* enhancer reporter, *brC12-lacZ* (magenta). Scale bar in (B), 100 μm. Box indicates the area of the wing pouch enlarged in (B’-B”). Scale bar in (B’), 50 μm. (C-C”) CtBP overexpression in *CtBP*^*Q229\**^ clones via the MARCM technique could fully suppress the upregulation of *brC12-lacZ* (use of the long isoform E is shown). Scale bar in (C), 100 μm. Box indicates the area of the wing pouch enlarged in (C’-C”). Scale bar in (C’), 50 μm. (D-F) Expression levels of *brC12-lacZ* were assessed in *yki*^*B5*^; *CtBP*^*Q229\**^ double-mutant clones. (D) A schematic showing the technique used to generate the double-mutant clones shown in (E, E’, F, F’). (D) *UbxFlp*-induced mitotic recombination independently generates GFP-negative *yki*^*B5*^ clones and RFP-negative *CtBP*^*Q229\**^ clones, resulting in four genetically-distinct cell populations: clones mutant for *yki* but wild-type for *CtBP* are red, clones mutant for *CtBP* but wild-type for *yki* are green, clones that have at least one functional copy of both *CtBP* and *yki* are yellow, and double-mutant clones lack all fluorescent markers and hence appear black. (E-F) Clones of all four genotypes could be recovered in a single disc, allowing for assessment of the *brC12-lacZ* reporter in each. Scale bar, 100 μm. (F-F’) A magnification of the dashed boxes in (E, E’) Compared to wild-type tissue (dotted outline), the *brC12-lacZ* reporter is increased in *CtBP* mutant tissue (double-asterisk) and decreased in *yki* mutant tissue (single-asterisk). Double-mutant tissue (solid outline) shows intermediate levels of the *brC12-lacZ* reporter.

### Immunostaining and Antibodies

Imaginal discs were dissected from wandering third instar larvae were fixed in PBS + 4% paraformaldehyde for 20 minutes at room temperature, washed and permeabilized in PBS + 0.1% Triton-X, and blocked in PBS + 0.1% Triton-X + 5% goat serum (G9023, Sigma) or donkey serum (D9663, Sigma). Primary antibody stains were performed overnight at 4°C, while secondary antibody stains were performed for 2-4 hours at room temperature.

Primary antibodies were diluted in the same solution used for the blocking step and used at the following concentrations: 1:50 goat anti-dCtBP (dN-20, Santa Cruz Biotechnology); 1:500 rabbit anti-cleaved DCP-1 (9578S, Cell Signaling Technology); 1:500 rabbit anti-β-Galactosidase (559762, MP Biomedicals); 1:500 mouse anti-β-Galactosidase (Sigma); 1:500 mouse anti-GFP; 1:500 rabbit anti-HA (C29F4, Cell Signaling Technology); 1:400 guinea pig anti-Yki; 1:1000 rabbit anti-dFos (a gift from D. Bohmann). Alexa Fluor-conjugated secondary antibodies were from Thermo Fisher Scientific and diluted 1:500 in the appropriate blocking solution. Nuclei were visualized with DAPI (1:1000). Tissues were mounted in SlowFade Diamond Antifade Mountant (Thermo Fisher Scientific). Confocal images were obtained on a Zeiss LSM 700 and processed using ImageJ.

### Imaging and processing of adult structures

Adult structures were imaged using a Leica transmitted light microscope (TL RCI, Germany). For adult wing experiments, wings were removed from adult females and mounted onto slides using Gary’s Magic Mounting Medium. The outline of each wing was traced using the Polygon tool beginning at the alar-costal break, and the bound area was quantified in ImageJ.

## Results

### CtBP is a negative regulator of tissue growth

In an unbiased mosaic screen for genes involved in growth regulation (described in (Tapon *et al.*, 2001)), we identified an allele of the gene *CtBP* that conferred a growth advantage to homozygous-mutant tissue. Homozygous mutant clones of this allele, *CtBP*^*A147T*^, generated by *eyFlp/FRT*-mediated mitotic recombination, were overrepresented in the adult eye when compared to clones of the parental *FRT82B* chromosome (**Figure 1A-B**). Conserved orthologs of *CtBP* are found in diverse taxa. It encodes a transcriptional co-repressor and was originally identified for its ability to bind the C-terminus of the adenovirus E1A oncoprotein in mammalian cells (Boyd *et al.*, 1993). Subsequent work has shown that, in both vertebrates, which have two CtBP genes, and invertebrates, which have just one, the CtBP proteins can form interactions with a large number of DNA-binding transcriptional repressors to regulate gene expression (reviewed in (Turner and Crossley, 2001; Chinnadurai, 2007)). Many of these repressors possess a PxDLS motif that is important for CtBP-binding. In *Drosophila*, the *CtBP* gene is predicted to produce multiple short (379-386 a.a.) and long (473-481 a.a.) protein isoforms via alternative splicing (**Figure 1E** **and** **Supplementary figure S1**).

The *CtBP*^*A147T*^ allele failed to complement the lethality of a previously described null allele of *CtBP* (*CtBP*^*87De-10*^, which we will refer to henceforth by its nonsense mutation *CtBP^Q229*^)*. This null allele causes embryonic lethality in homozygous animals, while animals homozygous for the *CtBP*^*A147T*^ allele survive to the late pupal stage, suggesting that the *CtBP*^*A147T*^ allele recovered in our screen is a hypomorph. It has been reported previously that *CtBP*^*Q229\**^ clones are larger than wild-type clones in mosaic eye imaginal discs (Hoang *et al.*, 2010). We observe this result as well (**Figure 1C**), with adult eyes consequently exhibiting large mutant clones. In addition to this published null allele (Poortinga *et al.*, 1998), we used CRISPR/Cas9-directed mutagenesis to generate an allele of *CtBP* with a 13-bp deletion at amino acid position 148 on the *FRT82B* chromosome. The deletion causes a frameshift mutation, *CtBP*^*N148fs*^, and consequent premature stop codon. Mutant clones of this likely null allele also display a growth advantage in mosaic eyes (**Figure 1D**).

Owing to its pleiotropic effects on gene expression, *Drosophila* CtBP has been implicated in a number of developmental processes, including embryonic segmentation (Nibu *et al.*, 1998; Poortinga *et al.*, 1998) and appendage patterning (Biryukova and Heitzler, 2008; Hoang *et al.*, 2010), yet less is known about its specific function in tissue growth control. While our results are consistent with a previous report’s findings that CtBP regulates tissue growth within the developing eye (Hoang *et al.*, 2010), its mechanistic basis is not known. To first determine whether CtBP also had similar functions in other imaginal discs, we generated CtBP mutant clones throughout the imaginal discs using *hsFlp/FRT*-mediated mitotic recombination. Because the *CtBP* mutation is flipped over an *FRT* chromosome bearing a fluorescent marker, we could directly compare the area of *CtBP* mutant tissue, marked by the absence of the fluorescent marker, to that of the wild-type twin-spot tissue. We found that for all three *CtBP* alleles, homozygous mutant clones comprised nearly twice as much of the whole wing disc as did twin-spot tissue, while in control mosaic discs, the total areas of unmarked and twin-spot tissue are roughly the same (**Figure 1F-H**). We also noticed that, particularly for the two null alleles, *CtBP*^*Q229*^ and *CtBPN148*^*fs*^, mutant clones had smooth edges when compared to wild-type clones (**Figure 1F, G**).

Overexpressing each individual protein isoform of CtBP in the wing pouch consistently led to an overall size reduction of the adult wing, as well as ectopic vein tissue, likely reflecting CtBP’s dual roles in growth and patterning (**Supplementary Figure S1, B-E**). However, the severity of these phenotypes was variable between isoforms and even when assessing the effects of a single isoform, particularly those that fell into the “short” category. In general, overexpression of the “long” isoforms resulted in smaller, rounder wings and mild ectopic vein formation while overexpression of the “short” isoforms reduced wing size while maintaining relatively a normal shape and in excessive ectopic veins (**Supplementary Figure S1 B-E**). Overall, the phenotypes associated with both *CtBP* loss-of-function and overexpression indicate that CtBP functions as a negative regulator of growth.

### CtBP limits tissue growth by repressing *bantam* expression

Given CtBP’s identity as a transcriptional co-repressor, we explored the possibility that CtBP may normally restrict tissue growth by regulating the expression of known growth promoters. We examined readouts of the Hippo pathway due to some similarities in phenotype to loss-of-function mutations in the Hippo pathway (mutant clones larger with smooth boundaries). We tested the effects of manipulating *CtBP* function on *in vivo* transcriptional reporters for the Yki target genes *bantam* (*ban*), *expanded* (*ex*), and *Diap1*, as well as the protein expression pattern of Yki. In *CtBP*^*Q229\**^ clones, expression of a *ban3-GFP* enhancer reporter (Matakatsu and Blair, 2012) was increased, particularly near clonal boundaries (**Figure 2A’-A”**). We also observed a slight upregulation of *ex* expression (**Supplemental Figure S2 A-A’’’**), as assessed by an *ex-lacZ* enhancer trap (*ex^697^* (Boedigheimer and Laughon, 1993)) though this seemed to occur mostly along the cells in contact with wild-type tissue. Expression of a *Diap1-GFP* transgenic reporter (Zhang *et al.*, 2008), however, was unchanged in *CtBP*^*Q229\**^ clones (**Supplemental Figure S2 B-B’’’**) suggesting that the effect was small and hence only detected with the more-sensitive readout. An antibody stain for Yki was not appreciably altered by loss of CtBP, showing, if anything, a slight decrease in overall levels within a *CtBP*^*Q229\**^ clone (**Supplemental Figure S2 C-C’’**).

The *ban* gene encodes a microRNA (miRNA) that promotes growth by stimulating cell proliferation and inhibiting apoptosis (Brennecke *et al.*, 2003). Consistent with the observed increase in *ban3-GFP* enhancer activity, a *GFP-ban* sensor, which inversely reports ban miRNA levels (Brennecke *et al.*, 2003) showed decreased GFP levels in *CtBP*^*Q229\**^ clones (**Figure 2B-B’’’**), thus indicating higher levels of *ban* miRNA activity. Conversely, FLP-out Gal4 clones that overexpress *CtBP* show higher *GFP-ban* sensor expression, indicating reduced *ban* miRNA levels (**Figure 2C-C’**). We note that this effect is shown via overexpression of the short CtBP.I-HA isoform (**Figure 2C-C’**), but we also tested the short CtBP.A-HA and long CtBP.E-HA isoforms and observed increased *GFP-ban* sensor activity indicative of reduced *ban* expression in clones overexpressing these isoforms as well (not shown).

These results suggest that increased *ban* expression may be responsible for the overgrowth phenotype of *CtBP* mutant tissue. To assess this possibility, we tested whether reducing *ban* levels can suppress the overgrowth of *CtBP* mutant clones using the MARCM technique, which allows for the expression of UAS transgenes within mosaic clones (Lee and Luo, 1999). Expression of a previously described *ban* sponge transgene (*ban-sp*), which contains repeated *ban*-binding sites to deplete the endogenous pool of available *ban* miRNA (Becam *et al.*, 2011), significantly reduced the growth of *CtBP*^*Q229\**^ MARCM clones (**Figure 2D-G**; compare **Figure 2G** **to** **2F**; quantified in **Figure 2H**). The average size of *CtBP*^*Q229\**^ clones expressing *ban-sp* did not differ significantly from that of control *FRT82B* clones expressing *ban-sp* (compare **Figure 2G** **to** **2E**; quantified in **Figure 2H**). Taken together, these results suggest a role for *CtBP* in regulating tissue growth via repression of the growth-promoting miRNA *ban*.

### *CtBP* regulates a minimal *bantam* enhancer, *brC12* via Yorkie-dependent and Yorkie-independent mechanisms

We next searched for factors that might mechanistically link CtBP to *ban* regulation. Within the *ban3* enhancer sequence used to generate the *ban3-GFP* reporter (Matakatsu and Blair, 2012) is an approximately 400 bp sequence, *brC12* (**Figure 3A**), that has been previously shown to function as a Yki-dependent minimal *ban* enhancer (Oh and Irvine, 2011). When we tested whether this enhancer is sensitive to changes in CtBP, we detected upregulation of a *brC12-lacZ* reporter gene in *CtBP*^*Q229\**^ mutant clones that were positively marked with GFP using MARCM (**Figure 3B-B’’**). Overexpressing *CtBP* in *CtBP*^*Q229\**^ MARCM clones could fully suppress this increased enhancer activity (we show use of the long isoform E in **Figure 3C-C’’** but also observed a similar effect using the short isoform A). By identifying a minimal sequence by which *CtBP* is capable of regulating *ban*, we sought to use this readout to further probe the mechanism linking *CtBP* to tissue growth.

Because the minimal *brC12* enhancer is known to be regulated in a Yki-dependent manner (Oh and Irvine, 2011), we next tested whether *yki* is required for the increased reporter activity caused by loss of *CtBP*. This was done by generating *yki*^*B5*^; *CtBP*^*Q229\**^ double-mutant clones using *UbxFlp* and double *FRT* chromosomes and assaying *brC12-lacZ* reporter activity (**Figure 3D-F’’**). Because the *yki* and *CtBP* alleles are carried on separate *FRT* chromosomes, they can be independently flipped over uniquely marked *FRT* chromosomes (**Figure 3D-F’’**). As outlined in **Figure 3D**, this will result in four genetically-distinct populations of cells distinguishable by their fluorescent markers: (1) cells that lack *yki* function but have at least one functional copy of *CtBP* are red; (2) cells that lack *CtBP* function but have at least one functional version of *yki* are green; (3) cells that have at least one functional copy of both *yki* and *CtBP* are yellow; and (4) cells that are double-mutant i.e. lack both *yki* and *CtBP* function also lack both fluorescent markers and hence appear black. Consistent with a previous report (Yu and Pan, 2018) and the results shown in **Figure 3E-F’**, *brC12-lacZ* reporter activity is completely abolished in *yki* single-mutant tissue (single-asterisk in **Figure 3E’**). Expression is elevated in *CtBP* single-mutant tissue (double-asterisk in **Figure 3E’**) as compared to wild-type tissue (dotted outline in **Figure 3D’, E’**). If the upregulation of *brC12* enhancer activity in *CtBP* mutant clones were solely due to excessive *yki* function, then double-mutant tissue would be predicted to have the same level of reporter activity as that found in *yki* single-mutant tissue. However, *brC12-lacZ* expression actually appeared slightly higher in the double-mutant tissue (solid outline in **Figure 3F-F’**) than in the *yki* single-mutant tissue (single-asterisk in **Figure 3F-F’**), but was still lower than that of the *CtBP* single-mutant tissue. This intermediate phenotype indicates that there is both a *yki*-dependent and *yki*-independent component by which *CtBP* regulates *ban* enhancer activity. Consistent with this observation, expression of *CtBP*-RNAi in *yki* null resulted in clones that were intermediate in size between *yki* null clones and clones expressing the *CtBP* RNAi alone (**Supplemental Figure S3**).

### *brC12* activity depends on a basal level of JNK signaling

We sought to further explore the Yki-independent component to *ban* regulation. We identified a putative AP-1 binding site, TGAGTCA (Angel *et al.*, 1987), within a highly conserved region of the *brC12* sequence; AP-1 is the key downstream transcriptional effector of the JNK pathway suggesting that *ban* enhancer activity may be regulated by JNK signaling. To test this, we assayed the effects of various JNK pathway manipulations on the *brC12-lacZ* reporter, using a *ptc-Gal4* driver. As shown in **Figure 4A**, *brC12-lacZ* is expressed relatively uniformly throughout the wing imaginal disc. Blocking JNK signaling via expression of a dominant-negative version of the *Drosophila* JNK protein Basket (Bsk-DN) (Adachi-Yamada *et al.*, 1999) completely abolished reporter expression within the *ptc-Gal4* stripe (**Figure 4B**). This suggests that *ban* enhancer activity requires a basal level of JNK signaling that is normally present throughout undamaged discs.

**Figure 4.**
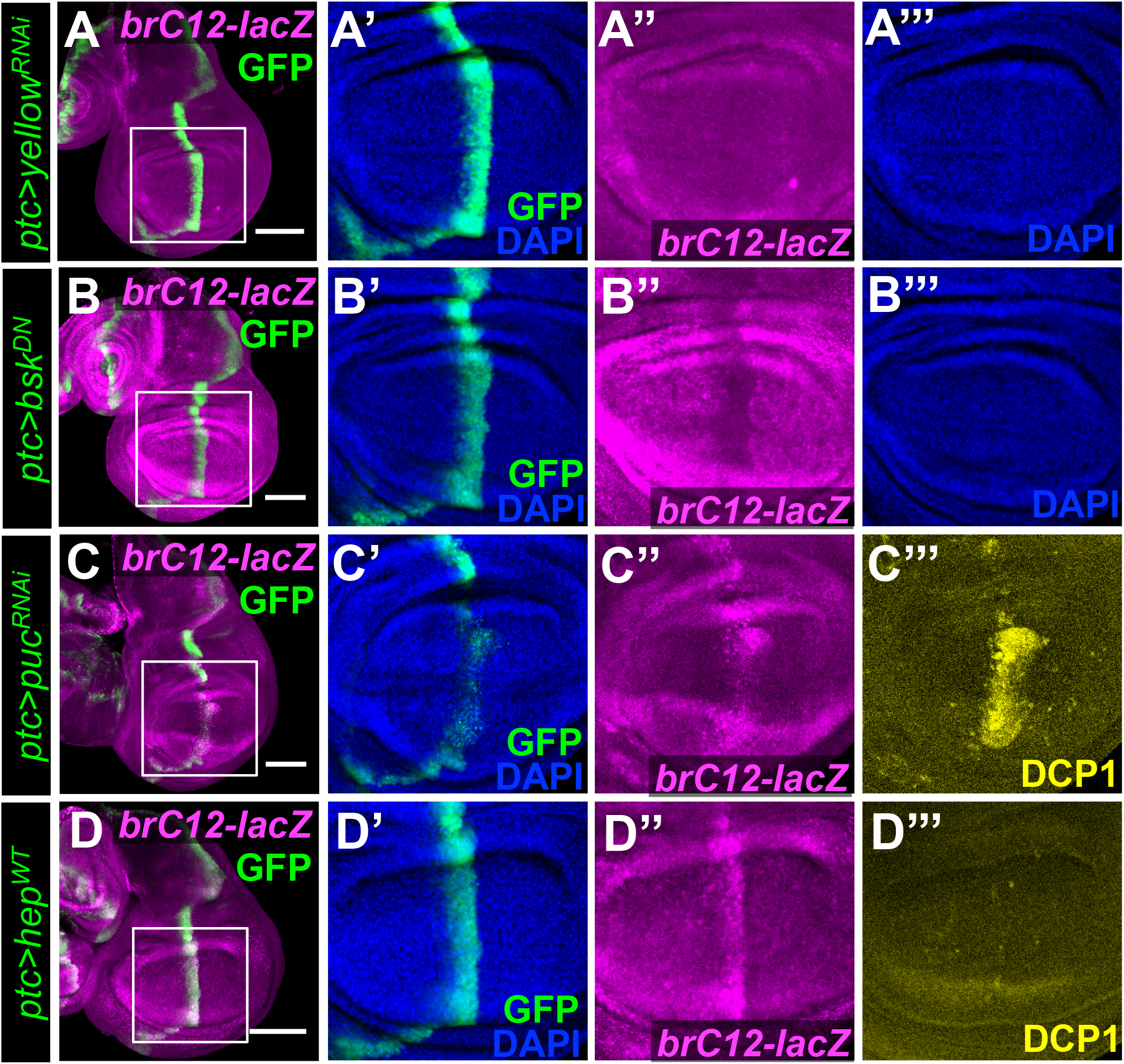
*brC12* activity is dependent on JNK signaling. (A-D) Manipulations of the JNK pathway were performed by expressing various UAS-transgenes in the *ptc*-domain (green) and the effects on the *brC12-lacZ* reporter (magenta) were assessed in wing imaginal discs. Boxes indicate the areas of the wing pouch that are enlarged in subsequent panels. DAPI shows nuclei (blue). (A-A’’’) A control disc showing *brC12-lacZ* reporter expression when *yellow* is knocked down. (B-B”’) Blocking JNK signaling using a dominant-negative form of Bsk/JNK reduces *brC12-lacZ* expression levels. (C-C”’) Strong pathway activation via knockdown of *puc* (C”) upregulates *brC12-lacZ* expression but also leads to apoptosis (C”’). (D-D”’) Elevating JNK pathway activity via expression of a wild-type form of *hep*/JNKK causes upregulation of *brC12-lacZ* (D”) without increasing apoptosis (D’’’). Scale bars, 100 μm.

Because the JNK pathway is known to promote apoptosis (reviewed by (Igaki, 2009)), this can complicate interpretation of *brC12-lacZ* expression changes under conditions of JNK pathway over-activation, as growth effectors are often upregulated during compensatory growth of lost tissue (reviewed in (Worley *et al.*, 2012)). For instance, elevating pathway activity via knockdown of the JNK phosphatase *puckered* (*puc*) resulted in an increase in both *brC12-lacZ* expression and apoptosis, as detected by DCP-1 staining (**Figure 4C**). However, when we mildly activated JNK signaling by overexpressing the JNK kinase Hemipterous (Hep) (Glise *et al.*, 1995), we observed elevated levels of *brC12-lacZ* expression without an appreciable increase in apoptosis (**Figure 4D**). Taken together, our results indicate that activity of the minimal *ban* enhancer is JNK-dependent and its activation by JNK can be uncoupled from JNK-induced apoptosis.

### CtBP also regulates *brC12* activity by antagonizing JNK signaling

Previously, while characterizing the role of *CtBP* in regulating regenerative plasticity, our group reported that loss of *CtBP* function results in the upregulation of several reporters of AP-1 activity (Worley *et al.*, 2018), suggesting that JNK and CtBP may have opposing effects on shared transcriptional targets (**Figure 5A**). Since we have demonstrated the minimal *ban* enhancer is activated by JNK signaling, we hypothesized that CtBP may normally antagonize this effect, which could explain why *brC12-lacZ* expression is increased in *CtBP* mutant tissue. To determine if CtBP regulates the *ban* enhancer in a JNK-dependent manner, we co-expressed Bsk-DN with *CtBP*-targeting sh*RNA* using *ptc-Gal4* and examined reporter expression. As described previously, shRNA-mediated knockdown can persist long after expression of the shRNA gene has ceased (Bosch *et al.*, 2016) (**Figure 5B**). Indeed, we observed persistent knockdown of the CtBP protein, as revealed by CtBP immunostaining, in cells that previously expressed, but stopped expressing Gal4 (**Figure 5C**). Notably, we consistently observed increased levels of the *brC12-lacZ* reporter outside of the current *ptc-Gal4* expression domain, indicated by *UAS-GFP* expression (marked by yellow brackets in (**Figure 5C-D**). Taking advantage of the persistent *CtBP* knockdown caused by *ptc-Gal4*-driven expression of the *CtBP-shRNA* transgene, we co-expressed the shorter-lived Bsk-DN protein, whose function is presumably limited to the current *ptc>GFP* domain, to compare *brC12* enhancer activity in *CtBP*-knockdown tissue in the presence or absence of JNK signaling (**Figure 5D**). Whereas *brC12-lacZ* levels were increased in cells lacking *CtBP* alone (marked by yellow brackets in **Figure 5D**), knocking down *CtBP* and blocking JNK signaling together brought reporter expression down to levels similar to what we observed in the non-Gal4/UAS-expressing tissue (**Figure 5D**). Thus, JNK signaling is required for the full upregulation of *brC12* activity caused by loss of *CtBP*. Notably, co-expression of Bsk-DN and *CtBP-shRNA* did not appear to reduce reporter levels to the same extent observed when overexpressing Bsk-DN alone (compare **Figure 5D’** to **Figure 4B”**), suggesting that there are JNK-independent mechanisms by which *CtBP* can regulate the *brC12* enhancer.

**Figure 5.**
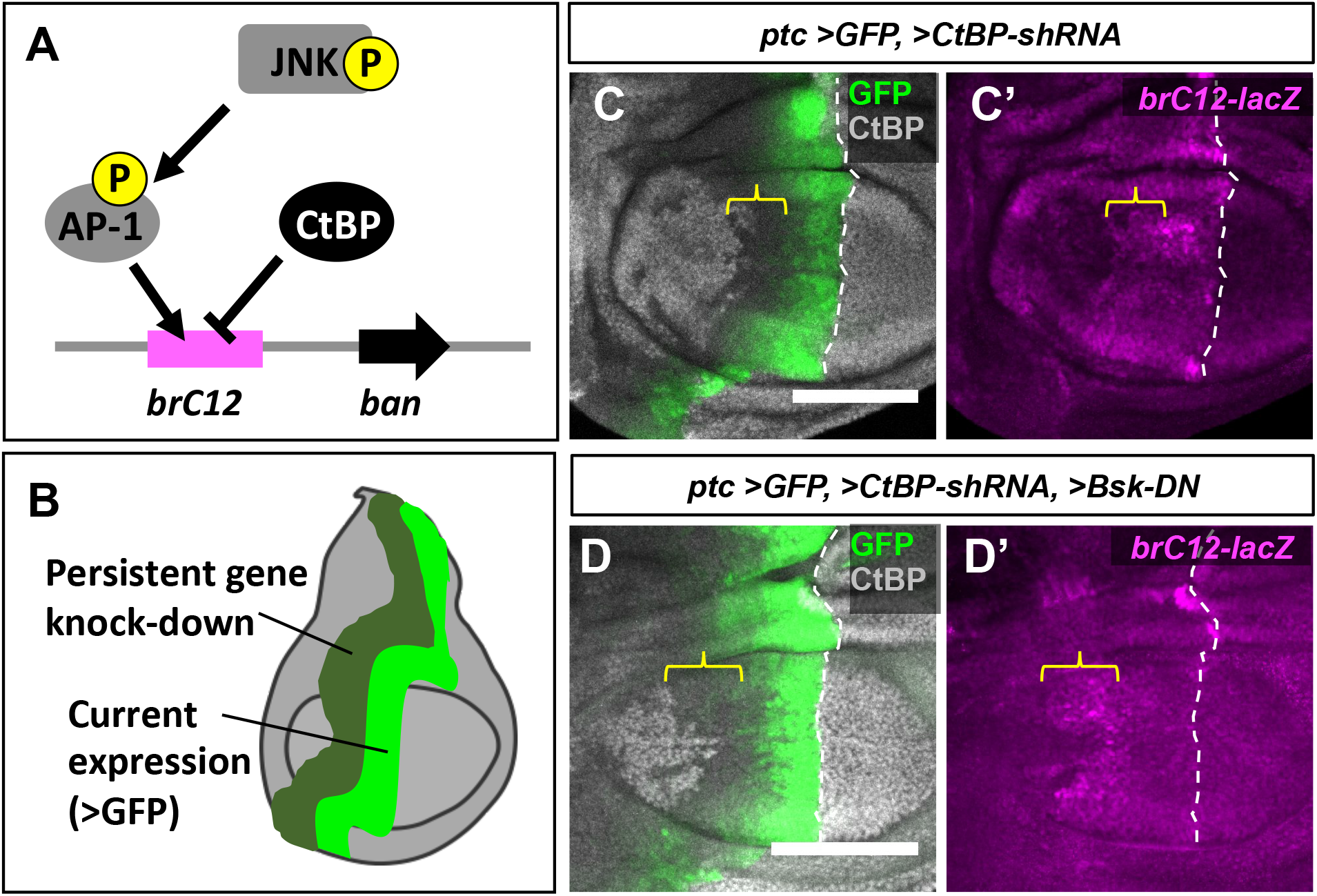
Blocking JNK signaling abrogates the increased *brC12* activity caused by loss of *CtBP*. (A) Model of *ban* gene regulation through the *brC12* enhancer. (B) Experimental design. (C) *CtBP-shRNA* expression using *ptc-Gal4* effectively knocks down *CtBP* in the *ptc*-domain (GFP) and results in prolonged CtBP knockdown in cells that expressed *ptc* earlier in development (yellow bracket). (C’) Upregulation of *brC12-lacZ* is observed in CtBP-knockdown tissue, including in tissue no longer expressing *ptc-Gal4* (yellow bracket). (D) Co-expression of a dominant-negative form of Bsk/JNK (D’) suppresses the *brC12-lacZ* upregulation caused by *CtBP* knockdown. Yellow bracket shows that in a region of the disc in which CtBP knockdown persists but JNK signaling is not blocked, the *brC12-lacZ* reporter remains elevated. All scale bars, 100 μm.

### Loss of distinct Fos isoforms leads to increased *brC12* enhancer activity

Given CtBP’s identity as a transcriptional co-repressor, one possible mechanism by which CtBP could antagonize JNK activity is through a functional interaction with AP-1. The AP-1 complex is typically thought of as a transcriptional activator, which, in *Drosophila*, is formed by heterodimeric interactions between the transcription factors Jun (also known as Jra) and Fos (also known as Kayak and Fra) (Perkins *et al.*, 1988; Riesgo-Escovar and Hafen, 1997); reviewed by (Alfonso-Gonzalez and Riesgo-Escovar, 2018). The *kayak* gene has five annotated isoforms. Intriguingly, protein sequence analysis reveals that 2 of the 5 predicted isoforms of Fos, isoform A and isoform G, possess the canonical CtBP binding motif, PxDLS (**Figure 6A**). Thus, we speculated that CtBP could be recruited to the *ban* locus by AP-1 complexes containing these Fos proteins to mediate transcriptional repression. If so, then eliminating these isoforms might phenocopy the effect of inactivating CtBP. To test this, we took advantage of a ~17kb chromosomal deletion, Df(3R)ED6315 which removes 3 of the 4 alternatively spliced 5’ exons of the *fos* gene and spans the transcriptional start sites for 3 of the 5 isoforms, including A and G (**Figure 6A**). When we generated clones that are homozygous for this deletion (**Figure 6B**), we observed elevated *brC12-lacZ* reporter levels within these clones (**Figure 6B”**) (note that *brC12-lacZ* expression is elevated in the adjacent dorsal hinge even in wild-type tissue). The chromosomal region that is removed by the Df(3R)ED6315 deficiency also contains two additional genes, *fos intronic gene* (*fig*) (**Figure 6A**), which encodes a predicted PP2C phosphatase (Hudson *et al.*, 2007), and the *lncRNA CR46110* (not shown), neither of which have well-characterized functions. While we cannot exclude the possibility that elimination of one or more of these factors may be causing the observed increase in enhancer activity, given the analysis presented thus far, we take this result to raise the possibility that some Fos isoforms may have a repressive function.

**Figure 6.**
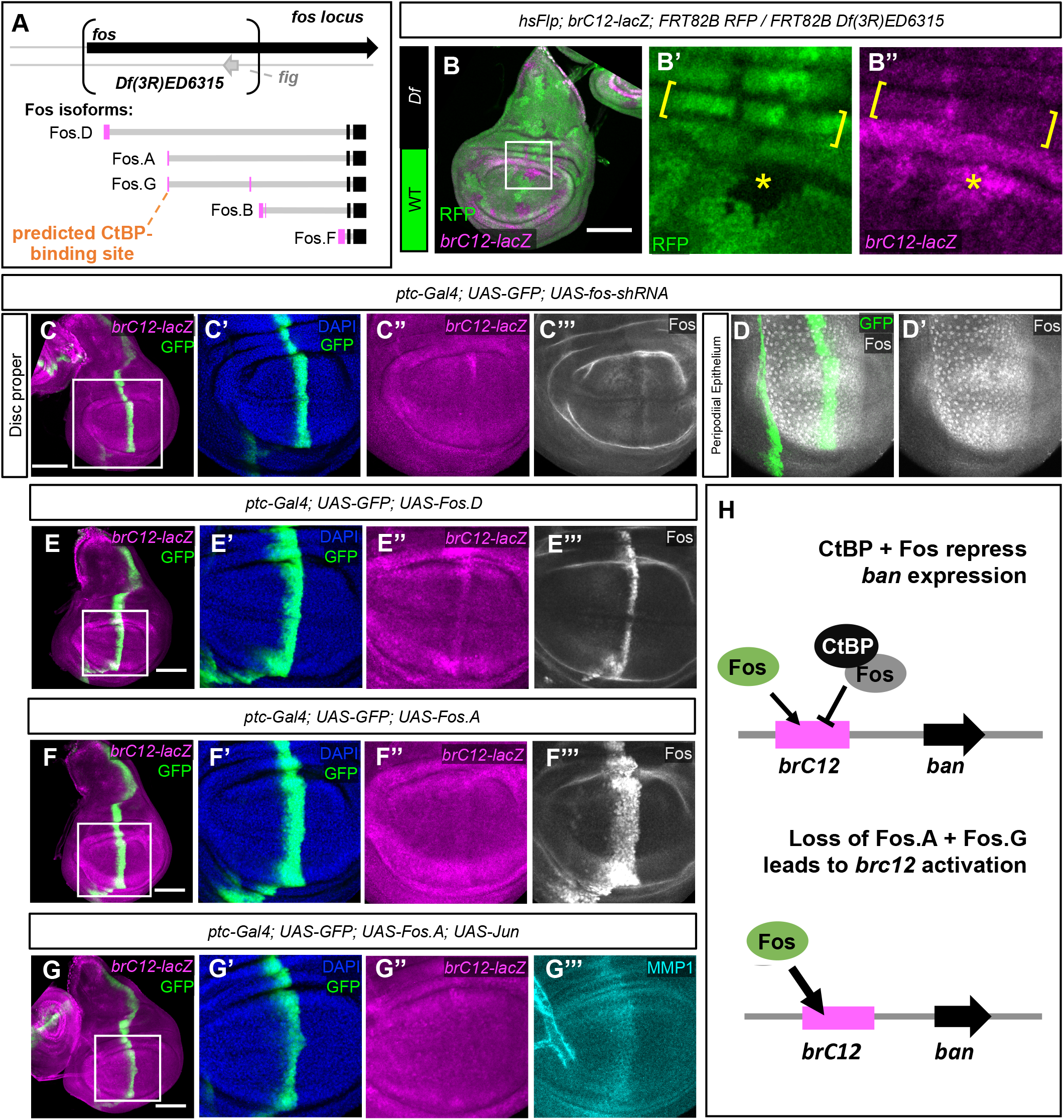
Loss of distinct Fos isoforms leads to *brC12-lacZ* upregulation. (A) A model of the *fos* genetic locus showing the 17-kb region removed by the *ED6315* deficiency (brackets) and five predicted protein isoforms. The black boxes represent coding exons shared by all isoforms while the magenta boxes represent coding exons present in only some isoforms. Intronic regions are represented by the horizontal grey lines; non-coding exons are not displayed. Isoforms A and G share identical first exons, containing a predicted CtBP-binding motif, PADLS. (B-B”) RFP-negative clones that are homozygous for the *ED6315* deletion show increased levels of *brC12-lacZ* (B”). Yellow brackets highlight hinge region with developmentally higher levels of *brC12-lacZ* expression. Asterisk markers clone with elevated *brC12-lacZ* expression. (C-G) Different *fos* manipulations were performed by expressing various UAS-transgenes in the *ptc*-domain (green) and the effects on the *brC12-lacZ* reporter (magenta) were assessed in wing imaginal discs. Boxes indicate the areas of the wing pouch that are enlarged in subsequent panels. DAPI shows nuclei (blue). (C) Knockdown of *fos* via expression of an shRNA predicted to target all isoforms does not show changes in *brC12-lacZ* reporter expression (C”). (C’’’) Fos protein levels (as detected by the Fos antibody) are low throughout the disc proper and reduced where the shRNA is expressed. (D-D’) The region of the same disc in C’-C”’ showing high Fos protein detection in the peripodial epithelium. (E-E”’) Overexpression of Fos.D, validated by Fos antibody staining (E”’), results in increased *brC12*-lacZ expression (E”). (F-F”’) Overexpression of Fos.A, validated by Fos antibody staining (F”’), does not affect *brC12-lacZ* expression (F”). (G-G”’) Co-expression of Fos.A and Jun still does not affect *brC12-lacZ* expression (G”) but does show increased levels of MMP1 protein (G’’’). All scale bars in figure, 100 μm. (H) Model of how CtBP could regulate the expression of *ban* through the Yki and JNK/AP-1 pathways.

To determine the overall effect of eliminating *fos* function, we knocked down *fos* using an shRNA that targets all isoforms (**Figure 6C**). Expressing this shRNA using *ptc-Gal4* had no appreciable effect on *brC12-lacZ* expression (**Figure 6C**). Immunostaining with an anti-Fos antibody confirmed Fos knockdown in the *ptc-Gal4* expression domain and also revealed that Fos is present at very low levels throughout the disc proper (**Figure 6C**), where most of our analyses of the *brC12-lacZ* reporter have so far been performed. Greater Fos protein expression was instead observed in the squamous portion of the peripodial epithelium (**Figure 6D**), where *ptc-Gal4* expression is limited to only a thin strip of cells just anterior to this domain.

The results presented are thus far consistent with three possible models. In one, Fos is not required for regulation of *brC12* enhancer activity, which is why *fos* knockdown had no effect on *brC12-lacZ* expression, and, despite the apparent AP-1 binding site identified in the minimal *ban* enhancer, JNK signaling activates the enhancer via a Fos/AP-1 independent mechanism. In this scenario, *brC12* reporter activity is sensitive to loss of some other factor removed by the Df(3R)ED6315 deficiency. A second model predicts that while Fos/AP-1 is normally involved in transcriptional activation, certain Fos isoforms may function, either primarily or in certain contexts, as transcriptional repressors. The result of knocking down all Fos isoforms could simply represent an “averaging” of these opposing effects. A third model could involve a hybrid of the first two models, in which JNK signaling activates the *ban* minimal enhancer via both AP-1-independent and AP-1-dependent mechanisms.

If distinct Fos isoforms do have specific effects on the *brC12* enhancer, then overexpressing individual isoforms could offer insights into their specific functions. We examined two isoforms, D and A, both of which are predicted to be affected by the Df(3R)ED6315 deficiency, but only one of which, isoform A, has the CtBP-binding motif PxDLS. Overexpression of isoform D (also known as the α isoform but which we will henceforth refer to as Fos.D) resulted in an upregulation of *brC12-lacZ* expression (**Figure 6E**), indicating that Fos.D is capable of activating the minimal enhancer. In contrast, overexpression of isoform A (also known as the β isoform but which we will henceforth refer to as Fos.A), which contains the PxDLS motif, did not upregulate *brC12-lacZ* expression (**Figure 6F**). Staining with an antibody that recognizes all Fos isoforms revealed that ectopic Fos.A was stably expressed to a greater degree, than ectopic Fos.D (**Figure 6F’’’, E’’’**). Because Fos and Jun together have been shown to form more stable protein-DNA complexes and thereby activate transcription to a greater extent than either of the proteins alone (Perkins *et al.*, 1990), we also co-expressed Fos.A with Jun (**Figure 6G**). This did not result in any appreciable change in *brC12* enhancer activity but was sufficient to upregulate a known target, *Mmp1* as assessed by MMP1 antibody staining (**Figure 6G’’’**). Thus, Fos.A does not appear to regulate the minimal *ban* enhancer under these conditions but is capable of increasing the expression of at least one other gene at least when overexpressed together with Jun. Thus it is possible that Fos.A activates transcription more weakly than Fos.D and that relative concentrations of specific isoforms of Fos, Jun and CtBP in individual cells could determine whether AP-1 functions as an activator or a repressor at specific promoters (**Figure 6H**).

## Discussion

We have shown that the transcriptional co-repressor *CtBP* functions to restrict growth, in significant part, via reducing the levels of the microRNA *ban*. The responsiveness of a well characterized *ban* enhancer, *brC12*, to changes in CtBP levels allowed us to identify the transcriptional regulators that are impacted by CtBP. We show that both Yki and AP-1 function at this enhancer and that *CtBP* can negatively regulate both of these inputs. In mammalian cells, YAP/TAZ and AP-1 have been shown to function synergistically at enhancers, especially to regulate genes that are involved in tumor progression and the acquisition of invasive characteristics. Our work raises the possibility that the mammalian orthologs of *CtBP*, *CTBP1* and *CTBP2* could function similarly to regulate the expression of genes regulated by those enhancers.

### Regulation of *ban* by *CtBP* via the Hippo pathway

We identified an allele of *CtBP* in a screen for mutations that enabled homozygous mutant clones generated during eye development to outgrow their wild-type sister clones. The observation that *CtBP* mutant clones are larger than wild-type clones has been previously reported (Hoang *et al.*, 2010), but the mechanism was not understood. Here we show that the overgrowth of *CtBP* mutant clones is characterized by and is dependent upon the increased expression of *ban*. We also show that the activity of a 410-bp *ban* enhancer is negatively regulated by CtBP. This enhancer has previously been shown to be activated by Yki. By generating clones that are mutant for either *CtBP* or *yki* and also double-mutant *yki* ^−^ *CtBP* ^−^ clones, we show that increased enhancer activity of *brC12* in *CtBP* clones has both Yki-dependent and Yki-independent components.

The mechanism by which Yki regulates *brC12* has been the subject of some debate. Because the *brC12* enhancer lacks a canonical Sd-binding motif, it was suggested that Yki could interact with the enhancer via a different DNA-binding protein such as Mad (Oh and Irvine, 2011). However, a requirement for *sd* was subsequently demonstrated by showing that Yki cannot activate *brc12* when *sd* function is removed (Yu and Pan, 2018). One explanation that would be consistent with both of these observations might be that the Yki/Sd complex does not bind to the enhancer directly but rather via a protein-protein interaction with a different DNA-binding protein such as Mad. Indeed, it has recently been shown that in mammalian cells, a complex containing YAP1 and the Sd ortholog TEAD4 can bind to enhancers of ecdysone-regulated genes not by interacting with DNA sequences, but rather via another protein, estrogen receptor α, that is already bound to the enhancer (Zhu *et al.*, 2019). Recent studies also suggest a mechanism by which CtBP can function as a repressor of Sd-regulated genes. CtBP can bind to the repressor Nerfin-1(Guo *et al.*, 2019), which in turn can interact with Sd (Vissers *et al.*, 2018).

### Regulation of *bantam* by the JNK/AP-1 pathway

While the JNK/AP-1 pathway is known to be activated in response to a variety of stress stimuli and has key roles in regeneration and tumorigenesis, its function in developmental growth is less understood. A role for the JNK pathway in regulating the growth of the wing disc is consistent with the original description of the phenotype of the *hemipterous* (*hep*) mutant where occasional viable escapers can lack wings; *hep* encodes a JNK kinase (JNKK) (Glise *et al.*, 1995). More recently, it has been reported that a localized stripe of phosphorylated (‘active’) JNK (pJNK) along the anterior-posterior (A/P) compartment boundary in the wing imaginal disc is critical for proper disc growth and that inhibiting this JNK activity results in a reduction in overall wing size (Willsey *et al.*, 2016). Despite using multiple anti-pJNK antibodies and a variety of staining conditions, we were unable to observe the reported pJNK stripe. However, our finding that the *brC12-lacZ* reporter can detect a seemingly uniform basal level of JNK signaling throughout the wing imaginal disc (**Figure 4B**) is consistent with the idea that low levels of JNK activity are important for normal growth. Changes in *brC12*-lacZ reporter expression elicited by reducing JNK signaling were not, however, detected with the *GFP-ban* sensor suggesting that the level of activation of this pathway could be very low.

We have identified a consensus AP-1 binding site within a conserved region of this enhancer sequence and shown that ectopic expression of the Fos.D isoform can activate the enhancer indicating that this enhancer can be activated by the canonical JNK/AP-1 pathway. However, we also have evidence that other *fos* isoforms (Fos.A) act differently and could even repress the *brC12* enhancer (**Figure 6B**). We observe that clones that lack a subset of Fos isoforms, including those with a predicted CtBP binding motif, activate the *brC12* enhancer, which suggests that the effects of Fos/AP-1 at this, and likely other enhancers, could be complex and depend on the relative levels of different isoforms including some that are capable of binding to CtBP (**Figure 6H**). The idea that CtBP, through its interactions with distinct Fos isoforms, may switch AP-1 from a transcriptional activator to a repressor is an exciting one, and could potentially be generalized to other examples of AP-1-dependent transcriptional regulation. In fact, a study of circadian rhythm behavior in *Drosophila* has previously demonstrated a unique requirement for Fos.D in regulating Clock (Clk) expression via its isoform-specific interaction with the Clk repressor VRI (Ling *et al.*, 2012). However, we also note that human *FOS* and paralogs *FOSB, FOSL1* and *FOSL2* lack a PXDLS motif. However, *FOS* does have a PxxDLS motif. It is also unclear how thoroughly the possibility of alternative isoforms has been explored at the *FOS* locus.

In addition to regulating the *brC12* enhancer via AP-1, JNK could conceivably also antagonize CtBP via the Yki-dependent branch of the pathway. JNK has been shown to increase Yki levels in the nucleus by phosphorylating Ajuba (Jub) which then inhibits Yki phosphorylation by Wts (Sun and Irvine, 2013). The effects of JNK on Yki seem context-dependent, however, since JNK can also inhibit Yki activity by more indirect mechanisms (Enomoto *et al.*, 2015).

### A potential role for CtBP as a tumor suppressor

We have now identified *CtBP* mutants in two different screens, one for mutations that increase plasticity during regeneration and another that detects mutations that increase tissue growth. Both of these findings hint at a possible role for mammalian *CtBP* orthologs in cancer since cancer cells typically display both of these characteristics. Chromatin profiling of mammalian cancer cell lines has shown that sites occupied by TEAD4 are often close to AP-1 binding sites and that the two pathways can function synergistically to activate target genes (Verfaillie *et al.*, 2015; Zanconato *et al.*, 2015; Liu *et al.*, 2016). Indeed many of the genes that seem to be regulated in this way seem to function in tumor progression and in promoting cell motility and invasiveness. Given the ability of *Drosophila* CtBP to regulate both Yki- and and AP-1-mediated gene expression it is possible that the mammalian CtBP proteins could also function at enhancers where these pathways converge. Although we have used an enhancer from the *ban* locus to study this phenomenon, since *ban* is not found in mammals, the relevant transcriptional targets must be different. Humans have two *CtBP* paralogs, *CTBP1* and *CTBP2*, which could potentially function with some level of redundancy. Increased expression of both genes has been observed in some cancers suggesting that they could function as oncogenes (Dcona *et al.*, 2017). However, CtBP has also been shown to antagonize some of the oncogenic properties of the adenovirus E1a protein (Subramanian *et al.*, 2013), suggesting that *CTBP1* and *CTBP2* could also function as tumor suppressors. Although mutations in these genes have been detected in human cancers as per data generated by the TCGA Research Network (https://www.cancer.gov/tcga), at least to our knowledge, the functional consequences of these mutations have not been evaluated. Assessing the impact of the mutated forms of *CTBP1* and *CTBP2* on YAP/TEAD and AP-1 transcription could be informative.

## Supplemental Figure Legends

**Figure S1.**
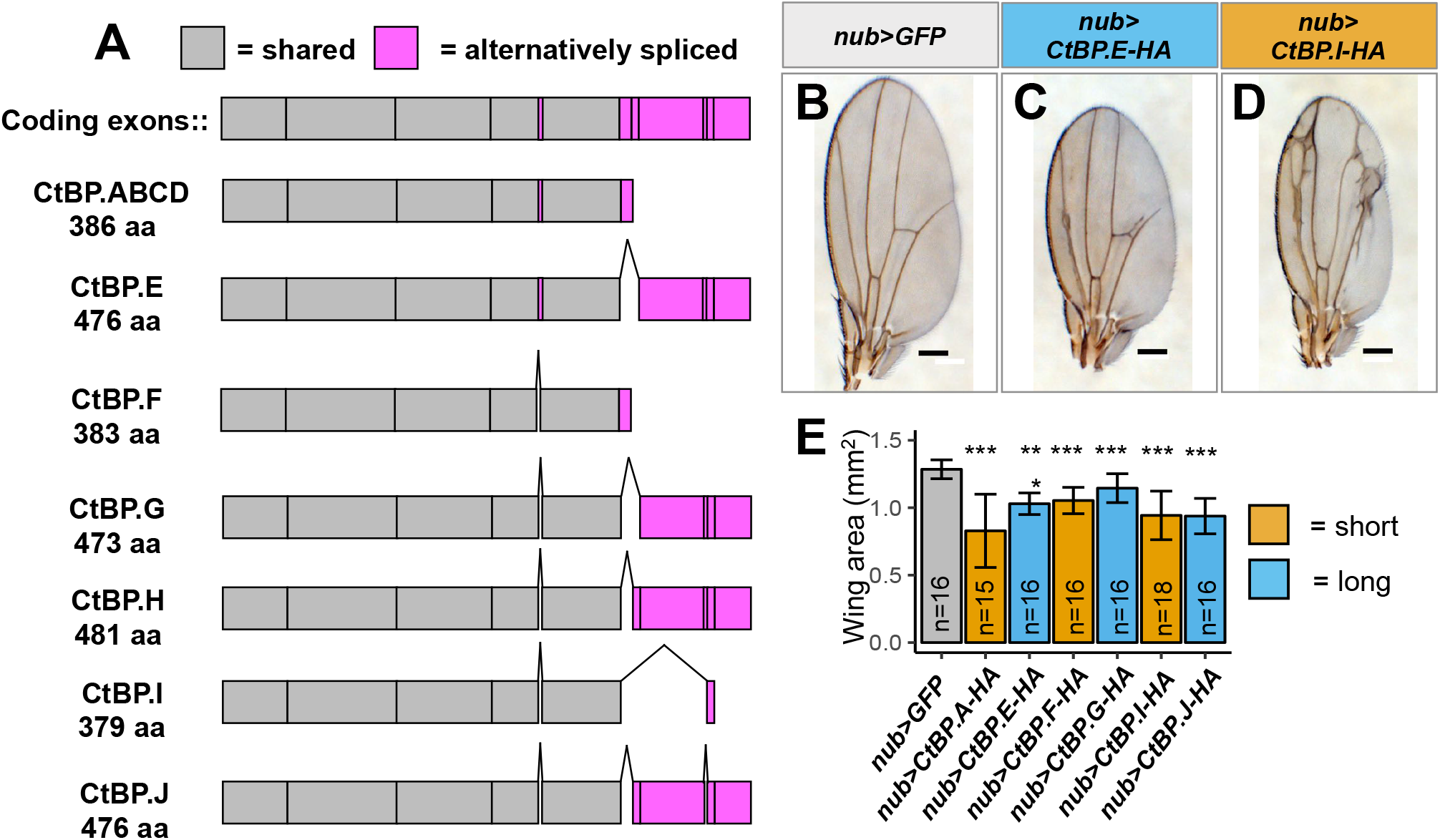
Isoforms of CtBP and effects of overexpression. (A) A schematic showing the distinct protein structure of the seven isoforms predicted to be encoded by the *CtBP* gene. The letters correspond to the encoding *CtBP* mRNA as referenced on FlyBase, with “ABCD” indicating four unique *CtBP* mRNAs with identical protein sequences. The green boxes indicate coding exons that are shared by all seven isoforms, while the magenta coding exons are alternatively spliced. (B-E) Overexpression of both short and long isoforms using *nub-Gal4* result in smaller adult wings. (B-D) Scale bars, 200 μm. (E) Quantification of wing area from overexpressing each CtBP isoform using *nub-Gal4.* n≥15 wings and data are presented as mean ± S.D. The individual isoform being overexpressed is ordered alphabetically, with short isoforms (379-386 a.a.) shown in orange and long isoforms (473-481 a.a.) shown in blue. Pairwise comparisons of the wing areas from the *nub>GFP* control and each isoform overexpression condition were performed using a student t-test and significance is displayed above each overexpression condition. \*\*\**P* ≤ 0.001.

**Figure S2.**
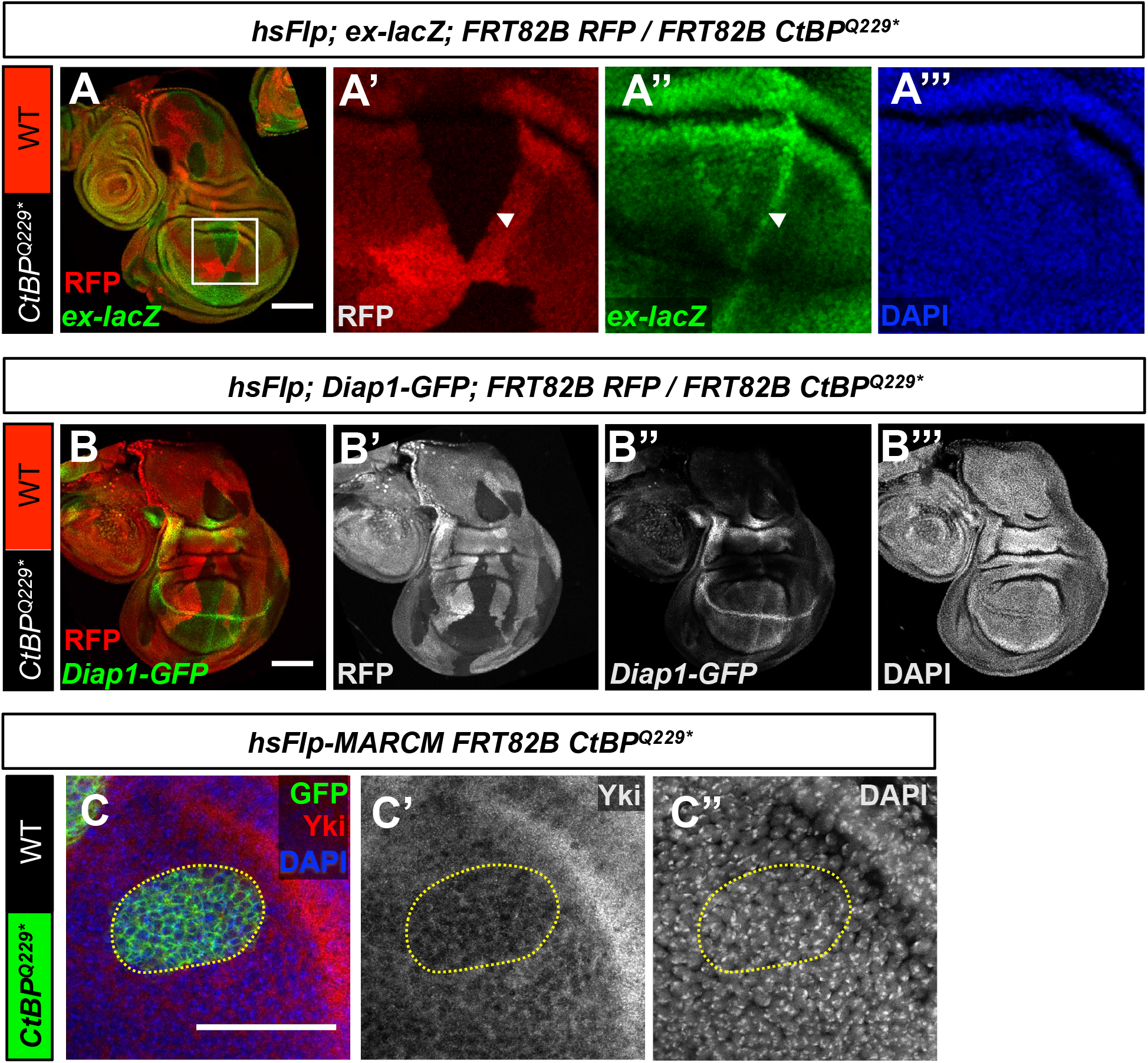
Additional analyses of Yorkie activity in *CtBP* mutant clones. (A-A’’’) *CtBP*^*Q229\**^ clones (unmarked) show increased levels of *ex-lacZ* (green) along clonal boundaries (arrowhead). Box indicates areas enlarged in A’-A’’’. Scale bar, 100 μm. (B-B’’’) *Diap1-GFP* expression is not appreciably altered in *CtBP*^*Q229\**^ clones. Scale bar, 100 μm. (C-C’’) An antibody stain for endogenous Yki shows that it is slightly decreased in a MARCM *CtBP*^*Q229\**^ clone (outlined). Scale bar, 50 μm. DAPI shows nuclei.

**Figure S3.**
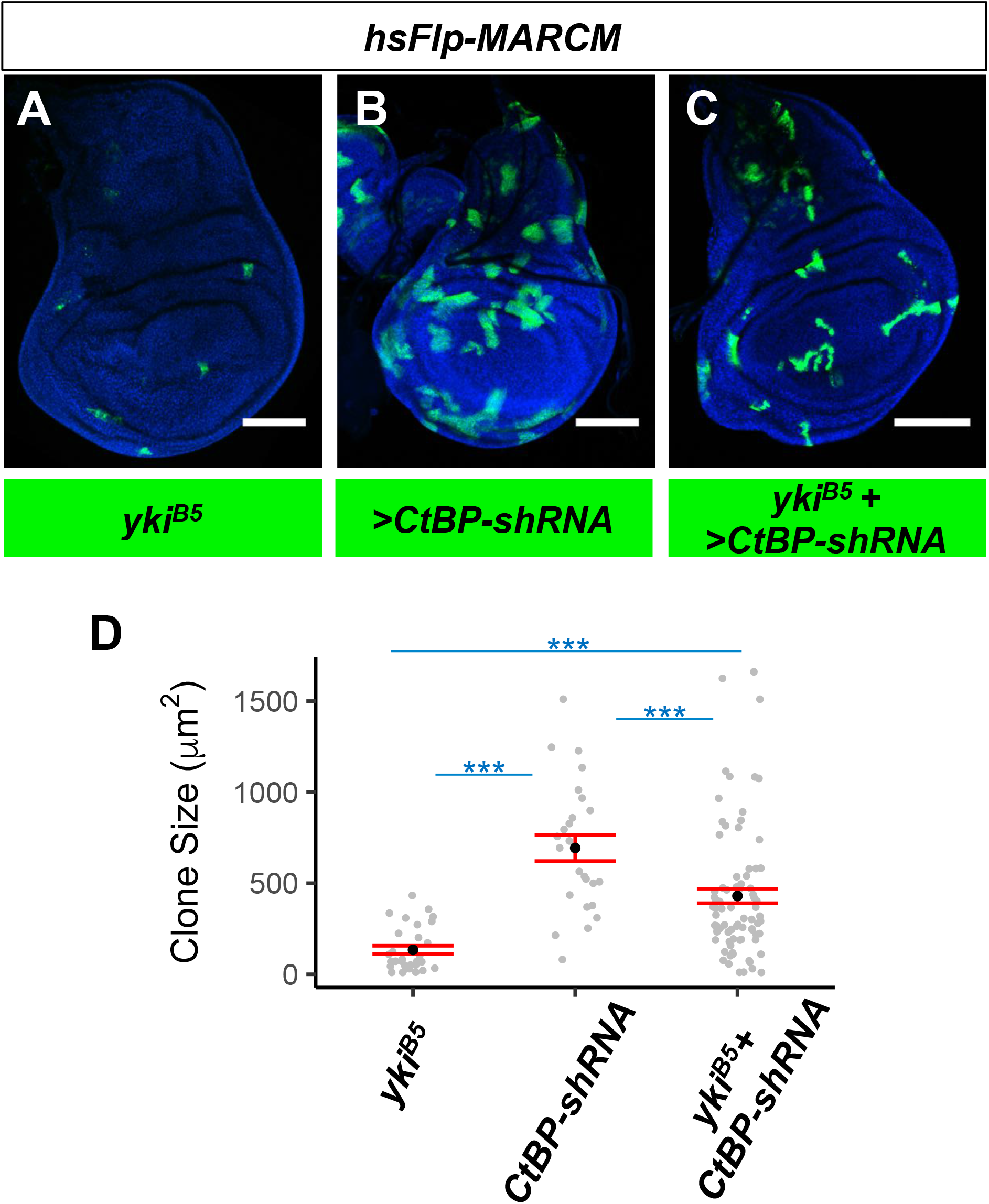
Loss of Yorkie partially abrogates the overgrowth caused by CtBP inactivation. (A-C) MARCM clone size assay. MARCM Gal4 clones (green) were induced by expression of *hsFlp* following a 1-hour heat-shock at ~48 hours after egg-lay. Discs were dissected ~72 hours after clone induction. Shown are representative wing pouches with (A) *yki*^*B5*^ mutant clones, (B) control clones overexpressing a shRNA targeting *CtBP*, and (C) *yki*^*B5*^ mutant clones overexpressing a shRNA targeting *CtBP*. (A-C) Scale bars, 100 μm. (D) Quantification of individual clone sizes (in pixels) from the genotypes shown in A (*n*=30), B (*n*=25), and C (*n*=80). Data are presented as mean ± S.E.M. Pairwise comparisons of each MARCM condition were performed using a Wilcoxon rank sum test. \*\*\**P* ≤ 0.001.

## Acknowledgments

We thank Seth Blair, Dirk Bohmann, Steve Cohen, Ken Irvine, Jin Jiang, Patrick Emery and Marco Milan for stocks or reagents. We thank Susan Wang, Kevin Gandhi and Larissa Alexander for technical assistance. We thank Kieran Harvey for discussions of unpublished results. IKH was funded by NIH grant R35 GM122490 and a Research Professor Award from the American Cancer Society (RP-16-238-06-COUN). TMS and JAB each received an NSF Graduate Research Fellowships and then a fellowship from the Cancer Research Coordinating Committee of the University of California. We thank the Bloomington Stock Center (NIH P40OD018537), Drosophila Genomics Resource Center, Developmental Studies Hybridoma Bank, and BestGene for stocks, reagents, and services.

